# Microbial community organization designates distinct pulmonary exacerbation types and predicts treatment outcome in cystic fibrosis

**DOI:** 10.1101/2023.07.21.550012

**Authors:** Stefanie Widder, Kristopher Opron, Lisa A. Carmody, Linda M. Kalikin, Lindsay J. Caverly, John J. LiPuma

## Abstract

Polymicrobial infection of the airways is a hallmark of obstructive lung diseases such as cystic fibrosis (CF), non-CF bronchiectasis, and chronic obstructive pulmonary disease (COPD). Intermittent pulmonary exacerbations (PEx) in these conditions are associated with lung function decline and higher mortality rates. An understanding of the microbial underpinnings of PEx is challenged by high inter-patient variability in airway microbial community profiles. We analyzed 880 near-daily CF sputum samples and developed non-standard microbiome descriptors to model community reorganization prior and during 18 PEx. We identified two communal microbial regimes with opposing ecology and dynamics. Whereas pathogen-governed dysbiosis showed hierarchical community organization and reduced diversity, anaerobic bloom dysbiosis displayed stochasticity and increased diversity. Microbiome organization modulated the relevance of pathogens and a simulation of antimicrobial treatment predicted better efficacy for hierarchically organized microbiota. This causal link between PEx, microbiome organization, and treatment success advances the development of personalized dysbiosis management in CF and, potentially, other obstructive lung diseases.

## Introduction

Obstructive lung diseases, such as cystic fibrosis (CF), non-CF bronchiectasis, and chronic obstructive pulmonary disease (COPD), are characterized by chronic polymicrobial bacterial infection of the airways. Intermittent increases in signs and symptoms of respiratory dysfunction, so-called pulmonary exacerbations (PEx), are associated with lung disease progression and mortality in these conditions.^1^ Despite their importance, the pathophysiologic events underlying PEx are unclear but generally believed to involve transient perturbation of host-microbial dynamics in the airways. Management of these events typically involves frequent, often aggressive, antibiotic treatment, which is intended to decrease bacterial burden and blunt host inflammatory response that contributes to lung pathology. This care carries considerable cost and treatment burden and is limited by drug toxicity and ever-increasing antimicrobial drug resistance.^2^ Thus, a better understanding of PEx remains a priority in efforts to improve care and enhance quality of life for persons with these conditions.^3^

The search, in cross-sectional studies, for common motifs in microbial community processes that drive PEx, particularly in CF, has been hampered by subject-specific microbiome configurations. Highly individual taxonomic profiles with context-dependent metabolic activities and signaling have been observed in numerous studies.^4–7^ Unpredictable and ill-defined onset of PEx, as well as personalized antimicrobial treatment schemes to manage PEx, further complicate analyses.^8^ A strategy that is capable of consolidating process communalities against the background of natural case variability is therefore required.^9^

Recent studies on microbial community networks have found that the precise configuration of dependencies among members defined their community role, as well as the dynamical behavior of the microbiome.^10–12^ Moreover, the formation of network clusters (i.e. the coexistence of microbial sub-communities) modulated robustness to external perturbation including antimicrobial therapy.^13, 14^ Interestingly, Palla et al. showed that temporal persistence of social clusters was facilitated by member turnover that depended on group size.^14^ The impact of community organization on pathogen virulence and on resilience to antimicrobial therapy in complex human microbiomes is understudied and remains largely unexplored for clinical applications.

In the present study, we developed non-standard descriptors that aggregate ecological and compositional properties of the CF lung microbiome and used these to identify PEx types with communal patterns. We then analyzed the organization of the CF microbiome in these backgrounds and revealed two fundamental dysbiosis states: a hierarchical community type controlled by the dominant pathogen and a stochastic type with blooming anaerobic taxa and high taxonomic turnover. Of note, the behavior of a focal pathogen was markedly different with different community hierarchy. Lastly, we modeled targeted antimicrobial treatment on data-inferred co-occurrence networks and observed that distinct community organizations significantly determined treatment outcomes.

## Results

### Compositional characterization of exacerbation trajectories

We aggregated a collection of 880 sputum samples from 11 people with CF that comprise 18 longitudinal sets, each spanning a period of clinical stability culminating with a PEx that prompted antibiotic treatment. More specifically, the collection included compositional microbiome profiling (based on bacterial 16S rRNA amplicon sequencing) of sputum samples collected near-daily from between 60 to one day(s) prior to the initiation of treatment; a mean of 49 sputum samples were analyzed per PEx. The time frame of 60 days prior to treatment was selected to accommodate potential early onset changes in the lung microbiome preceding symptom onset together with changes occurring during the acute PEx phase.

CF PEx are notoriously difficult to compare due to pronounced subject specificity of the lung microbiome, which typically overshadows potential communalities. Accordingly, we identified 1949 amplicon sequence variants (ASVs) among the 880 samples, with only eight ASVs present in every subject. For further analysis, we denoised the data set to 200 core ASVs by removing taxa present with an average relative abundance below 0.0075%. To quantify the degree of subject specificity in the data set, we performed a PERMANOVA test and calculated the effect sizes of clinical and demographic covariates on data variance. Covariates included subject, age, sex, clinical state (baseline health, exacerbation of symptoms^15^), *CFTR* mutation, and zygosity of the *CFTR* F508del allele. We found that individual subject is a strong predictor for ASV covariance (ω^2^ = 0.51, *p* < 0.001), followed by lesser effects of age (ω^2^ = 0.023, *p* < 0.001) and clinical state (ω^2^ = 0.003, *p* < 0.003). A summary of the data set is shown in **Table 1**, with inclusion criteria detailed in **Table S1**.

**Table 1.** Study cohort and sample characteristics.

| Study cohort |  |  |
| --- | --- | --- |
| Subjects |  | 11 |
| Age; mean (sd) |  | 35.3 (7.7) |
| Time series |  | 18 |
| by subject; mean (sd) |  | 1.6 (0.9) |
| Samples |  | 880 |
| by subject; mean (sd) |  | 80.0 (46.9) |
| by time series; mean (sd) |  | 48.9 (7.3) |
| by sex | female | 541 |
|  | male | 339 |
| by age group | <31 | 203 |
|  | 31-37 | 332 |
|  | 38-52 | 345 |
| by disease state | Baseline | 669 |
|  | Exacerbation | 211 |
| by CFTR mutation | F508del homozygous | 559 |
|  | F508del heterozygous | 214 |
|  | other | 107 |

### Identifying distinct PEx types using non-standard sample descriptors

To reduce subject-specific microbiome bias, we abandoned ASV composition as the *sole* sample descriptor. The CF lung microbiome is strongly conditioned by local oxygen gradients in accumulating mucus.^16–18^ Accordingly, we assembled all read counts in five higher-order groups reflecting the oxygen requirement of the 200 core ASVs (in the following called aerobicity groups^19^): strictly aerobic, strictly anaerobic, facultatively anaerobic, conventional CF pathogens (*Pseudomonas, Staphyolcoccus, Burkholderia, Haemophilus, Achromobacter* and *Stenotrophomonas*), and uncultivated taxa/unknown requirements. The conventional CF pathogens are generally considered aerobic but are capable of adaptation to hypoxic conditions. Building on aerobicity groups, we assembled the following non-standard descriptors for every sample: i) the ratio of CF pathogens to strict anaerobes, ii) the relative abundance of the most abundant CF pathogen, iii) the Shannon diversity index of the core ASVs, iv) the Chao1 richness of the core ASVs, and v) a community type classification using Dirichlet multinomial mixtures (DMM). The DMM model was implemented using aerobicity groups as input and identified six community types. Two community types were dominated by CF pathogens, three by anaerobes, and one by facultative anaerobic organisms. Community classes, selection of Dirichlet components and class distribution over cohort are presented in **Figure S1**.

We found that variance explained by the model (R^2^ = 0.62) was overall reduced compared to that based on the original ASVs (R^2^ = 0.78). Most importantly, effect sizes of subject bias decreased by 51% (**Figure 1A**). In the following, the data were ordinated using principal component analysis (**Figure 1B**), and the first three principal ordination components were used to group similar samples. We performed k-mer clustering and identified three distinct PEx clusters using χ^2^ statistics (**Figures 1C and D**).

**Figure 1.**
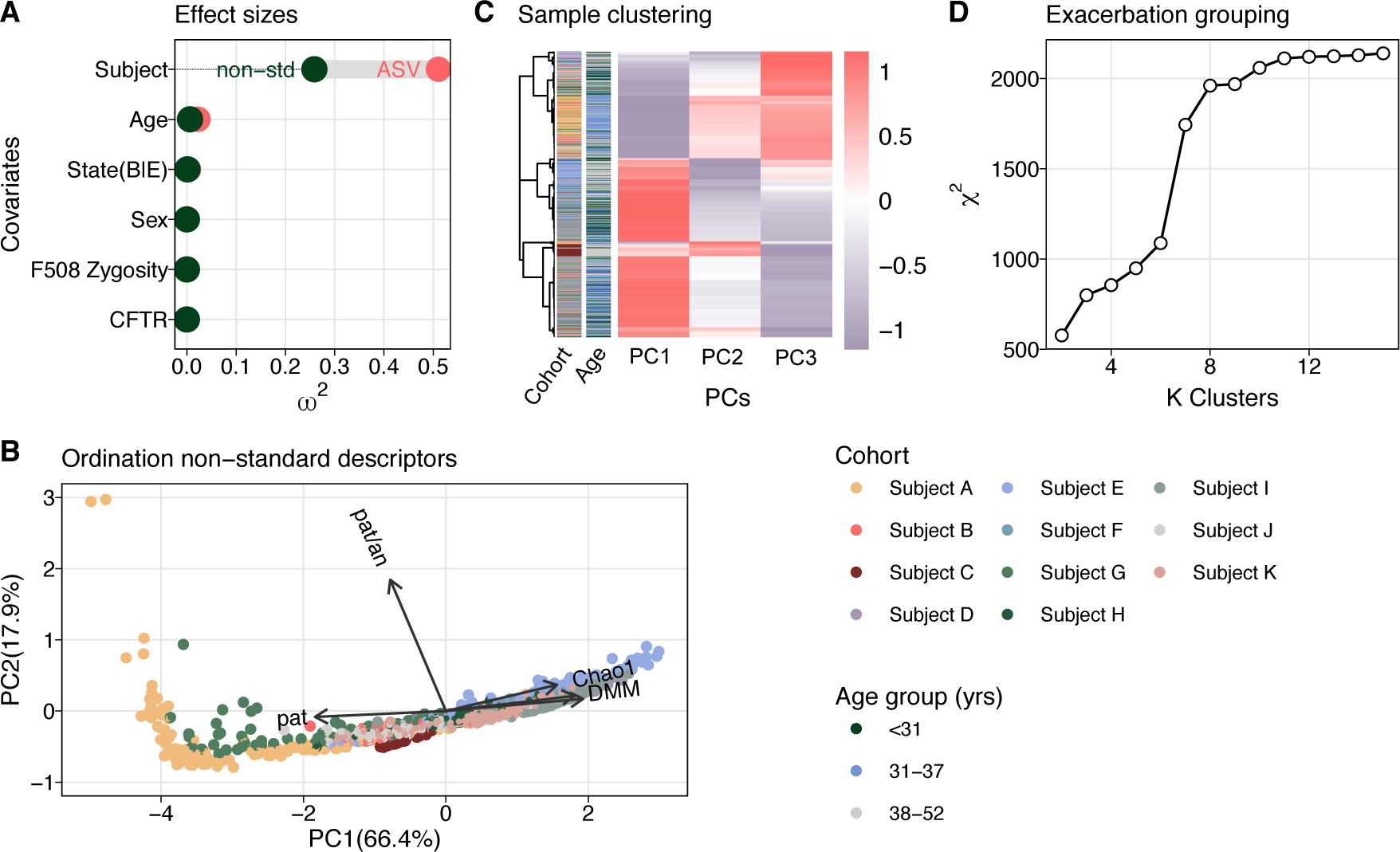
Stratification of similar exacerbations. **(A)** Covariate bias on explaining variance of microbiome data used in this study (n = 880 samples). Two PERMANOVA models contrasted the covariate effect sizes for ASV count data and non-standard sample descriptors. Partial values served as estimators of effect sizes. Covariates included subject ID, age group (< 31, 31-37, 38-52 years), clinical state (baseline, exacerbation), sex (female, male), F508del CFTR mutation zygosity (homozygous, heterozygous, n.a.), CFTR mutation (F508del^+/+^; 3 groups F508del^-/+^ and one other). **(B)** Data ordination using non-standard sample descriptors (principal component analysis, explained variance = 84.3%). Shannon diversity (Shannon), Chao1 richness (Chao1), relative abundance of the dominant pathogen (pat), ratio of counts derived from pathogens and anaerobes (pat/an) and sample classification by Dirichlet multinomial mixture model (DMM) served as model variables (n = 789 samples). Samples are colored by subject according to legend. **(C)** Sample-wise, hierarchical k-mer of ordinated data. Subject ID and age group, as well as Pearson correlation coefficients of the samples are depicted for additional information. Color code of cohort and age group according to legend. **(D)** Identification of optimal k-mer number by testing. The dependency between information gain and increasing cluster number is shown. First slope saturation served as a cutoff for the minimal number of clusters.

To classify subjects and their trajectories to a unique PEx cluster, we performed Spearman’s rank association (**Figures S2A and S2B**). After cluster assignment, the number of samples distributed as 286, 254, and 249, the number of subjects as 4, 3, and 4, and the number of PEx cycles as 6, 5, and 5 to PEx clusters 1, 2, and 3, respectively (**Figures S2C and S2D**). We found that sample association with cluster 1 was remarkably stable both at the level of individual PEx trajectories (60 days), as well as with subjects over time. On the contrary, more transition events were observed between clusters 2 and 3. Overall, subjects showed a tendency to persist either in cluster 1 or in clusters 2 or 3 despite recurrent acute treatment between time series included here (observation period of six months).

In summary, with the goal of revealing communal PEx types among subjects, we used aggregated measures of sample diversity, ecology and function that enabled us to reduce the organism-driven subject bias and group PEx trajectories with similar properties. We identified three clusters subsequently termed PEx types *PAT*, *AN1* and *AN2*, which displayed distinguishable microbiomes that were robust over half a year of observation time in CF lungs.

### Temporal behavior of microbiomes in distinct exacerbation regimes

We studied the time behavior of the three PEx types, aiming to elucidate the underlying ecological processes. We first asked whether PEx types could be characterized by classes of community configurations^20, 21^ and whether oxygen availability could motivate shifts in microbiome configurations.^22^ We assessed the distribution and temporal dynamics of DMM community types previously modeled from coarse-grained aerobicity groups (**Figures 2A, S1**). Unexpectedly, we observed no significant temporal dynamics of DMM communities within PEx types, indicating that the overall proportions of pathogens, anaerobes, facultative anaerobes and aerobes persisted over most of the PEx cycle with few exceptions. These sporadic shifts occurred only between comparable community classes, i.e. due to continuous transitions (increase or decrease) of taxonomic groups.

**Figure 2.**
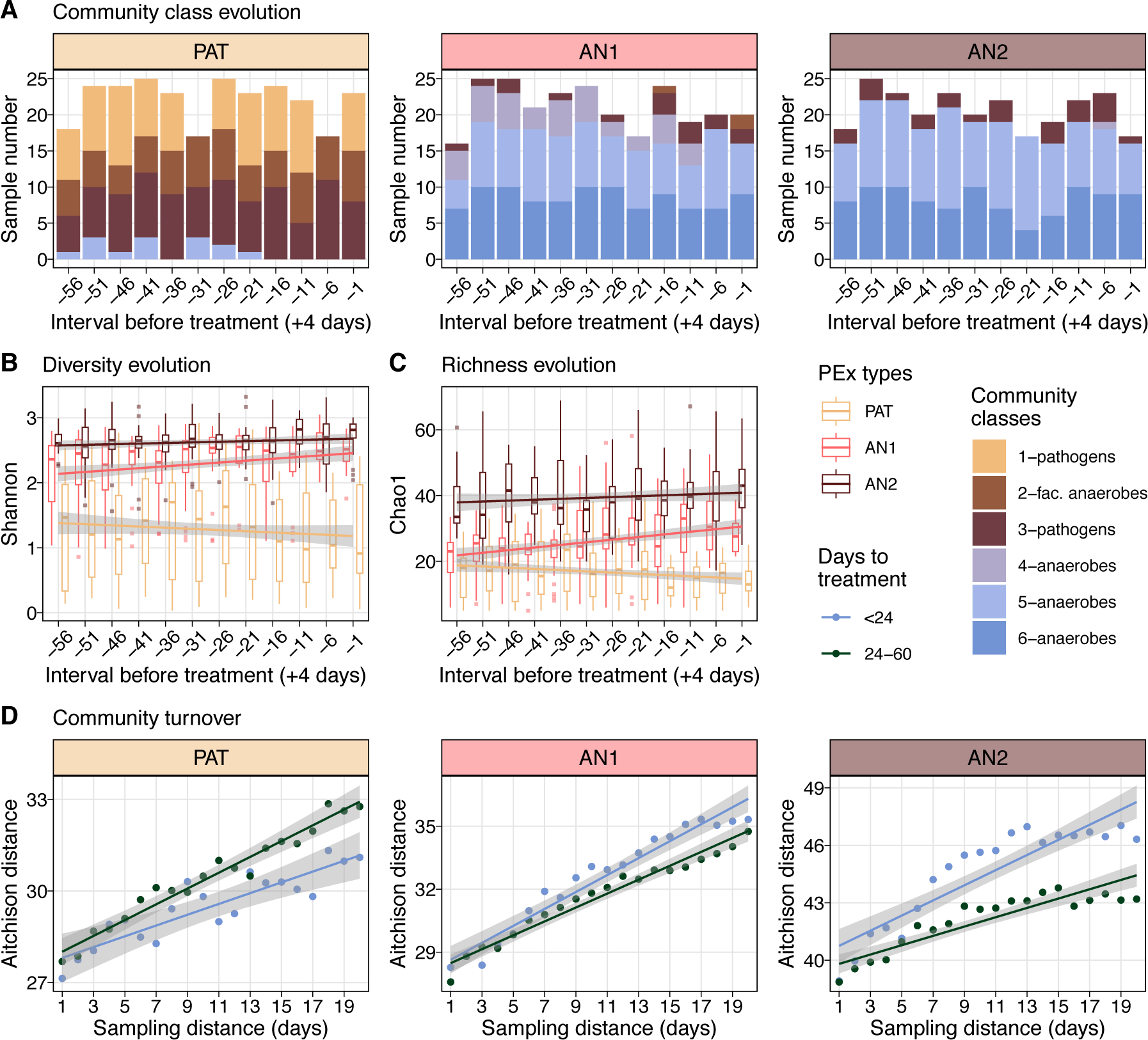
Temporal behavior of exacerbation types. **(A)** Time evolution of representative community compositions in three PEx types (PAT, AN1, AN2). Sample compositions were encoded by aerobicity groups and classified by a Dirichlet multinomial mixture (DMM) model into 6 compositional classes. Three classes were dominated by strictly anaerobic taxa, two by classical CF pathogens and one by facultative anaerobes (color coding according to legend). Community classes were independent of time towards acute treatment, but differed between PEx types (n = 789 samples). **(B)** Time evolution of microbiome diversity towards acute treatment. Microbiomes in all PEx types showed significant time dependency (LMM, *p*_*PAT*_ < 0.01, *p*_*AN*1_ < 0.05, *p*_*PAT*2_ < 0.05) with Shannon diversity decreasing in pathogen-dominated and increasing in anaerobe-dominated PEX types towards treatment. **(C)** Time evolution of community richness towards acute treatment. Chao1 richness changed significantly with time in pathogen-dominated and in one anaerobic PEx type (LMM,*p*_*PAT*_ < 0.001, *p*_*AN*1_ < 0.05). **(D)** Community turnover relative to the previous 20 days for three PEx types. Samples were grouped according to their relative collection term prior to acute treatment and mean values are plotted (in blue mean of samples collected < 24 days before treatment, in green 24-60 days). Aitchison distance from focal sample to n previously collected samples (sampling distance) is plotted to quantify community turnover. Turnover is significantly reduced in pathogen-dominated microbiomes and significantly increased in anaerobe-dominated microbiomes in a 23-days-interval before acute treatment (ANCOVA, *p* < 0.001 for each pairwise test).

Next, we analyzed the distribution of DMM communities between PEx types. Cluster 1 (*PAT*) comprised communities dominated by conventional CF pathogens, including *Pseudomonas*, *Burkholderia*, *Achromobacter*, *Haemophilus, Staphylococcus*, and *Stenotrophomonas*. PEx clusters 2 (*AN1*) and 3 (*AN2*), on the other hand, were driven by three distinct anaerobic community configurations. These results suggested that species sorting occurred in patients’ lungs according to oxygen requirements.^23^ Importantly, PEx proceeded in both aerobic and anaerobic communities.

Several recent studies investigated microbiome structure and rearrangement prior to PEx with inconsistent results.^24–27^ In short, neither pathogen load nor other individual organisms were consistent predictors for imminent PEx across larger patient cohorts. Here, we used the identified PEx types and analyzed diversity and richness over time in trajectories with similar microbiome properties. We implemented mixed effect models that tested time dependencies of Shannon and Chao1 for the three PEx types and corrected for confounders (subject id, PEx cycle id) (**Figures 2B and 2C**). All PEx types displayed significant diversity evolution across the baseline and PEx samples culminating in antibiotic treatment (p_PAT_ = 0.00126, p_AN1_ = 0.04462, p_AN2_ = 0.03672). The analogue analysis for Chao1 identified *PAT* and *AN1* to exhibit significant time dependency with time (p_PAT_ = 4.333e-05, p_AN1_ = 0.025).

Interestingly, richness and diversity decreased towards treatment for the pathogen-dominated communities (*PAT*) and increased for anaerobic clusters (*AN1* and *AN2*). Furthermore, it is important to note that the time dependency of richness and diversity were consistently small and ranged between η^2^ = {0.02, 0.06} for all PEx types. In short, we revealed that diversity evolves opposingly, as the microbial communities approached acute treatment. Of note, despite the modest effect size, these results have the potential to explain the inconclusive reports of previous studies that were conducted without consideration of PEx regimes.

### Species turnover displays antagonistic patterns in pathogen or anaerobe communities

Evidence suggests that changes in airway microbial community structures may precede the onset of clinical symptoms of PEx by days or even weeks.^7, 25, 28^ To determine the most likely time frame for such changes, we grouped samples by collection time (days before the initiation of treatment for PEx) and tested for significant differences of Shannon diversity (**Figure S3**). A split into 1-23 and 24-60 days before acute treatment showed statistically significant relative changes in all three PEx types.

We analyzed species turnover during onset of (0-23 days prior to treatment) and prior to (24-60 days) PEx by evaluating Aitchison distance between any two samples collected in an interval of one to 20 days in a subject-wise manner (**Figure 2D**). Overall, Aitchison distance was lower in the pathogen-dominated PEx type, displaying increasingly reduced turnover during the 23 days prior to treatment compared to 24-60 days prior (ANCOVA p < 0.001). Interestingly, the anaerobic PEx types again exhibited antagonistic change patterns, with increased species turnover shortly before acute treatment (ANCOVA p < 0.001 for both tests).

Together, the previous results suggested two PEx regimes with antagonistic temporal behavior.

### Different community organization characterizes pulmonary dysbiosis

The detailed organization of interactions and dependencies throughout an ecological community predefines its emergent, dynamical capabilities.^29^ In particular, resilience to perturbations such as antimicrobial treatment, community robustness and the stabilizing effect of keystone organisms were previously attributed to properties of dependency networks.^10, 11, 14^ Therefore, it is not only important to identify the most relevant CF pathogen in the microbiome, but to understand how the background community organization impacts the focal driver organism, modulates its virulence and contributes to stability.

We inferred 589 co-occurrence networks in a sliding window of 20 samples along each time series and analyzed network topologies by PEx type (n_PAT_ = 222, n_AN1_ = 192, n_AN2_ = 175, detailed description in Table S2).

In **Figure 3**, we studied the topology of the giant components, which are the largest connected portions of the network and hence the most impactful for microbiome dynamics. For PEx type *PAT*, we observed a reduced number of organisms and associations in the giant component, as well as increased betweenness centrality (Wilcoxon p < 0.001 for each pairwise test, **Figures 3A, 3B and 3C**) in contrast to PEx types *AN1 and AN2*. Graph betweenness centrality measures the extent of centralized organization reinforcing effective communication patterns.^30^ In analogy to interaction networks, we examined network hierarchy of the microbial co-occurrence networks across the PEx types.^31^ We inferred degree distributions p(k) together with clustering distributions p(C_i_) and compared the slope of a power law fit across all co-occurrence networks within the same PEx type (**Figures 3D and 3E**). While the fit to degree distributions displayed similar slopes (α_*PAT*_ = −0.71, α_*AN*1_ = −0.6, α_*AN*2_ = 0. −69, *R*_*PAT*_ = −0.64, *R*_*AN*1_ = −0.67, *R*_*AN*2_ = −0.73, *p* < 0.001), the slope of the node clustering distribution differed significantly between PEx types (α_*PAT*_ = −0.91, α_*AN*1_ = −0.27, α_*AN*2_ = −0.28). The pathogen-driven PEx type *PAT* showed the strongest descent of clustering probability by more than 3-fold, indicating a steep microbiome hierarchy. These compelling results supported the hypothesis of a pronounced, hierarchical dysbiosis. Moreover, the anaerobic PEx types *AN1* and *AN2* also exhibited weaker correlation between fits and data (*R*_*PAT*_ = −0.61, *R*_*AN*1_ = −0.3, *R*_*AN*2_ = −0.29, *p* < 0.001) suggesting flat organization and unpronounced community structure.

**Figure 3.**
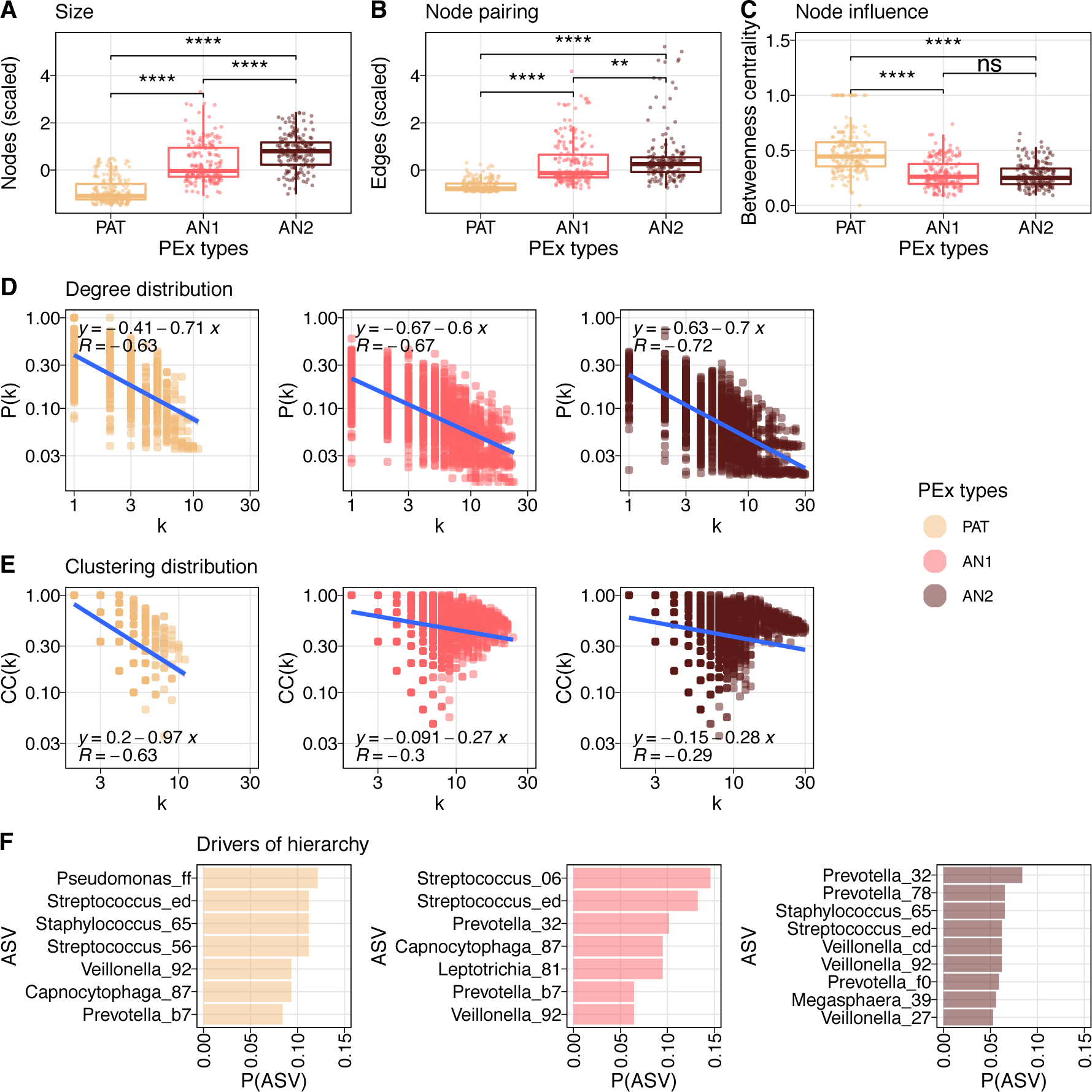
Microbial community organization in pulmonary dysbiosis. 589 co-occurrence networks were inferred trajectory-wise with SparCC. Graph properties of the giant component are depicted..**(A)** Boxplots of community sizes for three PEx types (PAT1, AN1, AN2). Pathogen-dominated communities were significantly smaller than microbiomes dominated by anaerobes (Wilcoxon, *p* < 0.001 for each pairwise test). **(B)** Boxplots of edge numbers for three PEX types. AN1 and AN2 displayed significantly more co-occurrences than PAT (Wilcoxon, *p* < 0.001 for each pairwise test). **(C)** Node influence on information flow within the microbiome. Betweenness centrality of every member in the giant component is presented for three PEx types. Information flow in PAT microbiomes was significantly higher than in AN1 or AN2 communities (Wilcoxon, *p* < 0.01 for each pairwise test). **(D)** Degree distribution over all co-occurrence networks in respective PEx type (color code according to legend). Probability of finding a network with degree k is plotted in logarithmic scale. Linear regression line is depicted with model parameters, correlation of regression with data is reported. Power law in all three PEx types with comparable slopes *s* = {−0.71, −0.7, −0.6}. **(E)** Clustering distribution over all co-occurrence networks in respective PEx type (color code according to legend). The distribution of clustering coefficients by network degree is plotted in logarithmic scale. Linear regression line is depicted with model parameters, correlation of regression with data is reported. Clustering for nodes with higher degree declined 3.5 times faster in pathogen-dominated organization than in anaerobe communities (*s* = −0.97, −0.28, −0.27, respectively). **(F)** Organisms residing in most hierarchical community positions. Top hierarchical positions were defined as nodes with highest 10% of all node degrees and lowest 10% of all clustering coefficients in the respective graph. ASV ranking in these positions (minimal appearance *P* = 0.05) is shown for three PEx types (color code according to legend).

To confirm that these results were driven by PEx types rather than sample diversity, we implemented independent linear mixed effect models for every graph readout, corrected for subject id and calculated effect sizes of PEx types and covariates Shannon diversity and Chao1 richness. We found that betweenness centrality, clustering, number of vertices and number edges significantly depended on PEx types with effect sizes being 2.9 times, 5.5 times, 8.9 times and 3.4 larger than the most effective diversity measures, respectively (*p*_*between*_ < 0.004, *p_cluster_* < 0.048, *p*_#*edge*_ < 2.7*e* − 7, *p*_#*edge*_ < 7.1*e* − 11). Clustering and betweenness centrality were statistically independent of tested diversity measures, while both diversities influenced edge numbers and richness graph size to a minor extent (**Figure S4**).

Next, we investigated which organisms preferentially occupied the most hierarchical positions of the communities and therefore likely controlled the overall microbiome dynamics. For each network, we identified the most hierarchical nodes (k > 90%, C_i_ < 10%) and assessed ASV frequencies on these positions (**Figures 3F**). In the hierarchical dysbiosis, *Pseudomona*s, *Staphylococcus* and *Streptococcus* were not only masters in hierarchy, but also belonged to the most abundant aerobicity group in the samples. In *AN2*, we observed a stronger variation of taxa in the most hierarchical nodes. Of note, one third of the ranking was occupied by various ASVs belonging to the genus *Prevotella* in these communities. We concluded that in anaerobic communities, individual species were less relevant for overall microbiome dynamics, and microbiome organization was increasingly stochastic and less well picked up by co-occurrence analysis. To the contrary, in pathogen dysbiosis, the hierarchies were conserved, occupied by few key organisms and well supported by a simple, centralized community organization.

### CF pathogens drive PEx dynamics in hierarchical, but not in flat community organization

The insight that bacterial organization appeared markedly distinct in the identified PEx regimes raised the important question whether microbiome organization could modulate pathogen importance or interfere with treatment outcomes in a foreseeable manner. As a first step, we used page rank as a statistical descriptor for network importance and compared the importance of conventional pathogens, strictly anaerobic, facultatively anaerobic, and strictly aerobic taxa in the community. We found that CF pathogens were differentially important for the CF community, with pathogens displaying significantly higher page rank in hierarchic than in flat community organization (**Figure 4A**). Next, we assessed pathogen dynamics in different community organizations using time series information. Previously, we demonstrated that stochastic ecological processes can be distinguished from interaction-driven processes by Fourier spectra inferred from the abundance changes of microbiota.^32^ Here, we determined the spectrum of every ASV per time series and inferred their noise color. We observed that pathogens exhibited interaction-driven dynamics (pink noise) in steep hierarchies only (**Figure 4B**). In flat anaerobe-dominated hierarchies, pathogens instead showed stochastic behavior (white noise, *AN2*) or anaerobic taxa dominated community dynamics (*AN1*). These results supported the hypothesis that CF pathogen activity depended on the community background.

**Figure 4.**
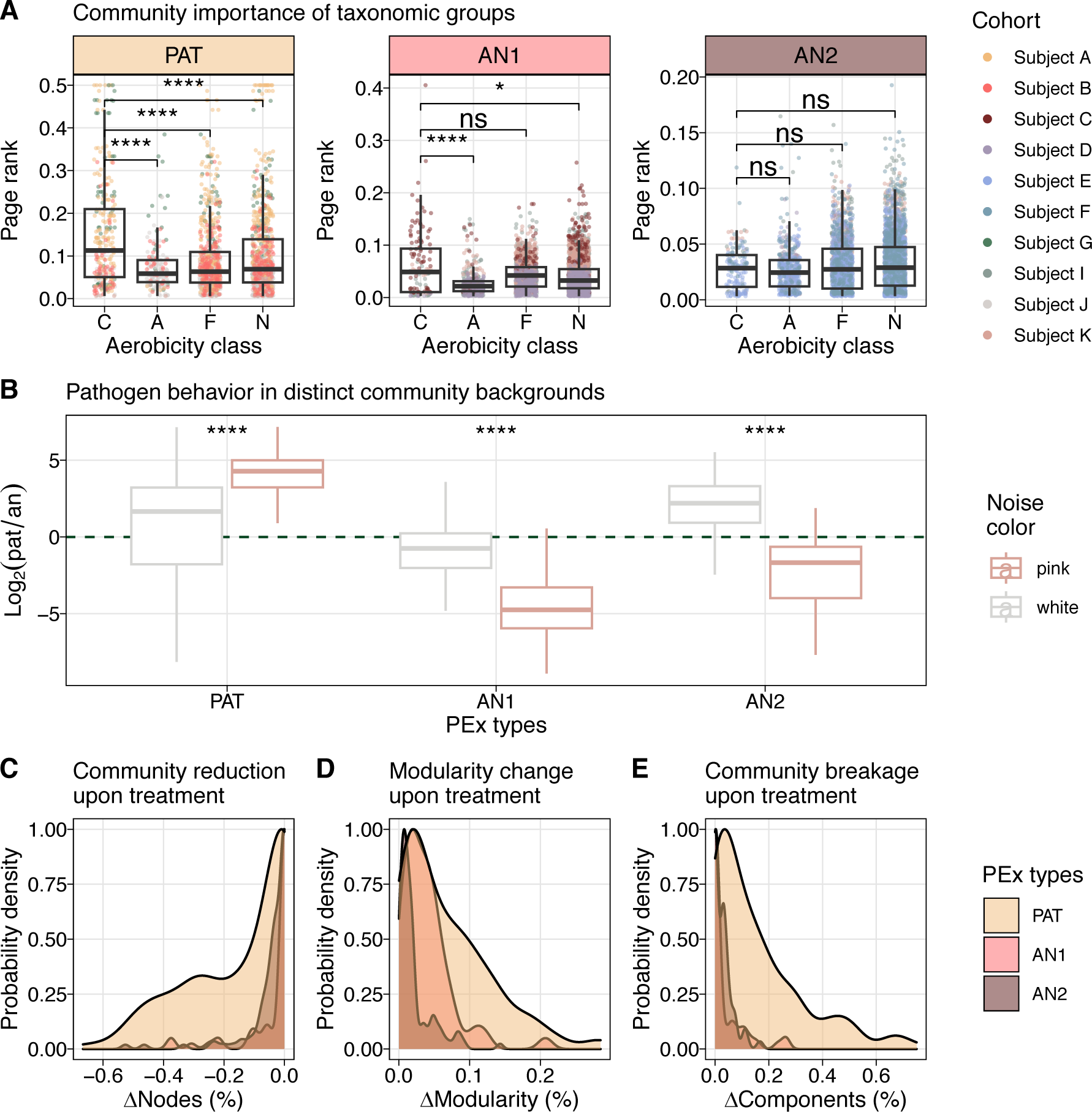
Community organization modulates treatment success and defines the importance of CF pathogens. **(A)** Community importance of organisms with identical aerobicity classes (C CF pathogens, A strict aerobes, F facultative anaerobes, N, strict anaerobes). Page rank of nodes in co-occurrence networks for three PEx types (PAT, AN1, AN2) are presented (n = 589 networks). Pathogens are significantly more important than any other aerobicity group in pathogen-dominated microbiomes, aerobic community members are significantly less important in AN1 (Wilcoxon, p < 0.0001 for all significant pairwise tests). No significant difference among organisms with distinct oxygen requirements were detected in AN2. **(B)** Log2 change of noise colors generated by pathogenic and anaerobic organisms in three PEx types. Noise color was assessed for every ASV by time series, 424 significant ASV classifications were obtained. The log2-ratio of pathogens to anaerobes (pat/an) associated with white or pink dynamics is plotted, green dashed line depicts shift of driver organisms (pathogens if *y* > 0, anaerobes if *y* < 0). White noise is the repercussion of stochastic time dynamics, pink noise the result of self-organized processes. CF pathogen displayed (self-)organized behavior in PAT only (*y*_STCU_ > 0), and furthermore showed stochastic behavior (*y*_*white*_ > 0, AN2) or were irrelevant (*y_white_* < 0, *y_pink_* < 0, AN1) in anaerobic communities. **(C-E)** Simulations of targeted pathogen removal from data derived co-occurrence networks. Most abundant pathogen in the sample was removed from the giant component and resulting community perturbation was assessed (n = 312 networks). Probability densities of the effects are presented. **(C)** depicts the relative size reduction (number of nodes) of the giant component after pathogen removal. **(D)** shows the relative increase of modularity after treatment and **(E)** presents the breaking of the giant component into smaller, unconnected components. Hierarchical community organization in PAT is significantly stronger disrupted by focal treatment, than flat organization in AN1 and AN2 (Kolmogorov-Smirnov, *p* < 0.01 for each pairwise test).

In previous graph theoretical work, it was demonstrated that dynamical fluctuation in response to external damage, as well as predictability of community response causally depended on the local network architecture of single layered and multiplex networks.^33–35^

In an analogue, simplified approach, we modeled the response of our empirical co-occurrence networks to focal pathogen removal by antibiotic treatment. We hypothesized that community organization affected the degree of network disruption and consequently community dynamics and likely treatment outcomes. We removed the most abundant pathogen from the major component of 313 networks (*PAT* = 152, *AN1* = 53, *AN2* = 108) and modeled network disruption by assessing modularity change, breakup into subcomponents, and size reduction after node removal (**Figures 4C, 4D and 4E**). We found that pathogen elimination from steeper background hierarchies resulted in significantly stronger topology disruption supported by all three topological parameters (Kolmogorov-Smirnov statistic in **Table S3**). We observed stronger change in modularity, increased disruption into unconnected subcomponents, as well as more pronounced size loss of the biggest connected component. We concluded that relevance of CF pathogens for microbial community dynamics and, by extension, likely also clinical course, was crucially shaped by community organization of the CF microbiome. Moreover, targeted treatment of pathogens resulted in distinct responses as a function of microbiome hierarchy. This model prediction suggested different clinical outcomes for treatment of the identical focal bacterium in dysbiotic or flat community background.

## Discussion

In this study, we harnessed the power of time series data to uncover similarities in PEx episodes in people with CF hosting individual microbiome profiles. Using a functional/ecological coarse graining enabled us to study both temporal and organizational aspects of CF lung microbial communities and their relevance for therapy outcomes. We identified two PEx regimes with distinct community structure, function and dynamics. One regime was shaped by the dominance of canonical CF pathogens, reduced biodiversity and decreased organismic turnover, indicating a pathogen-driven dysbiosis, while a second regime was characterized by a flaring of anaerobes. Moreover, community organization was markedly hierarchical in pathogen dysbiosis, yet increasingly stochastic and flat in anaerobic blooms.

To contextualize our observations, we examined these identified configurations in contrast to microbiota in lung homeostasis. In healthy lungs, the pulmonary microbiome shows neutral community dynamics.^36–38^ After Hubbell, diversity and abundance distributions of neutral communities can be explained by stochastic immigration and extinction events alone.^39^ To the contrary, in chronic lung disease, microbial interactions, local replication, and environmental adaptation become key for diversity and community dynamics, and the impact of dispersal diminishes.^5, 36^ Consequently, if neutrality is a property of microbial eubiosis in the lung, then microbial interactions can be considered a hallmark of pulmonary dysbiosis. Together with adaptation such interactions promote outgrowth of certain taxa to high relative abundances. Accordingly, we propose that both observed community states resemble different kinds of dysbiosis: The first, a structured, interacting community under the governance of an abundant, conventional CF pathogen, and the second, a globally successful functional guild that gains abundance by adapting to selective environmental pressures. Of note, similar community archetypes characterized by i) species-sorting or ii) mass effects were described in metacommunity theory, a framework for ecological community assembly and dynamics.^40^ The transition between the two metacommunity archetypes were explained by changes in dispersal due to altered spatial arrangements.^23^ Here, we speculate that subject-specific mucus accumulation and decreasing oxygen availability in the lung microenvironment determine CF dysbiosis states in equivalent ways.

Importantly, both community states are robust maladaptations to the disease conditions of the lung, which raises the question whether negative loops exist in the system that enable their dynamical stability.^5, 41^ We hypothesize that in the first state, functional adaptations of the dominant pathogens together with antimicrobial defense against microbial competitors provide important negative feedback, whereas limitations of available niche space stabilize the second regime.

The virulence of pathogenic bacteria depends on the biochemistry of the microenvironment among other factors. For example, *Pseudomonas aeruginosa* tightly regulates biofilm formation, as well as the production of siderophores and exotoxins to iron availability and oxygen levels.^42, 43^ Moreover, the fermentation products 2,3-butanediol and lactic acid produced by anaerobic members of the CF microbiome were reported to trigger quorum-sensing and further virulence.^44, 45^ Conversely, synergistic interactions such as metabolic cross-feeding affects pathogen growth and lowers the tolerance of *P. aeruginosa* to antimicrobial treatment independent of intrinsic antibiotic resistance profiles.^46, 47^ To clarify the relevance of community organization for medication strategies, we asked whether community organization can influence pathogen importance and modulate treatment outcomes. We modeled the focal depletion of the most abundant pathogen in empirical co-occurrence networks and recognized that distinct background communities shaped the outcome of the simulated antimicrobial treatment. Indeed, hierarchical communities were strongly affected, resulting in increased community reduction, while flat communities only responded weakly to simulated pathogen treatment. These results indicated that the influence of a CF pathogen, as well as its likely virulence and treatability systematically depended on its local network neighborhood. In graph theory, it was demonstrated repeatedly that the topology of interaction networks was intimately linked to its dynamical robustness.^33–35^ Interestingly, organization of microbial interactions has also been identified as a key element for robust composition forecasting.^48^ In complex networks, node organization not only determined the importance of hubs for systems dynamics^12^, but also controllability of dynamics supporting our previous conclusions.^49^

This study on the lung microbiome in CF showcases the importance of community organization for understanding microbiome dynamics in homeostatic and pathogenic human microbiomes. We identified two archetypes of dysbiosis in the lung - driven by pathogens or anaerobes - characterized their community structure and discussed stabilizing factors in the context of current graph and ecological theory. It is important to realize that the identified dysbiosis features can cancel out if analyzed cross-sectionally due to their antagonistic nature. This might explain the difficulties with establishing robust predictors for PEx thus far.^3^

We discovered that the relevance of pathogenic taxa for microbiome dynamics, disease progress and clinical treatability is clearly linked to the organization of the background microbiome. Our insights contribute to building theory for targeted dysbiosis management, antibiotic stewardship and personalized medicine in the age of the microbiome.

## Materials and Methods

### Study cohort and sample collection

283 of 880 sputum samples and DNA sequences included in this study have been reported previously (NCBI BioProject PRJNA520924). Expectorated sputum was collected from a cohort of people with CF as part of a long-term prospective study of CF airway microbiota. This study was approved by the Institutional Review Board of the University of Michigan Medical School (HUM00037056), and informed written consent was obtained from all participants. Subjects collected daily sputum samples at home, which were stored at either 4°C or -20°C before shipment to the University of Michigan for immediate storage at -80°C. Electronic medical records were reviewed for subject demographic and clinical data.

### Sample processing

Sputum samples were thawed on ice and homogenized with 10% Sputolysin (MilliporeSigma, Burlington, MA, USA). Samples were treated with bacterial lysis buffer (Roche Diagnostics Corp., Indianapolis, IN, USA), lysozyme (MilliporeSigma), and lysostaphin (MilliporeSigma) as previously described^50^, followed by mechanical disruption by glass bead beating and digestion in proteinase K (Qiagen Sciences, Germantown, MD, USA). DNA was extracted and purified using the MagNA Pure nucleic acid purification platform (Roche Diagnostics Corp., Indianapolis, IN, USA) according to the manufacturer’s protocol.

### Sequencing protocol and taxonomic annotation

DNA libraries were prepared by the University of Michigan Microbial Systems Molecular Biology Laboratory as described previously.^51^ In brief, the V4 region of the bacterial 16S rRNA gene was amplified using touchdown PCR with barcoded dual-index primers. Touchdown PCR was performed consisting of 2 min at 95°C, followed by 20 cycles of 95°C for 20 sec, 60°C (starting from 60°C, the annealing temperature decreased 0.3°C each cycle) for 15 sec, and 72°C for 5 min, followed by 20 cycles of 95°C for 20 sec, 55°C for 15 sec, and 72°C for 5 min and a final 72°C for 10 min. The resulting amplicon libraries were normalized and sequenced on an Illumina sequencing platform using a MiSeq Reagent Kit V2 (Illumina Inc., San Diego, CA, USA). The final load concentration was 4.0-5.5 pM with a 15% PhiX spike to add diversity.

Annotation was performed using the *dada2* pipeline in R.^52^ The data set was denoised removing all ASVs with < 0.0075 % average abundance across all samples as previously described^53^ and subsequently rarified using R package *vegan*.^54^ Sequencing data, taxonomic information and clinical meta data were organized in *phyloseq* objects for further analysis.^55^

### Community typing with Dirichlet multinomial mixture models

To stratify representative community classes across patients, Dirchlet multinomial mixtures (DMM) were inferred employing aerobicity classes as taxonomic level.^21^ Counts from ASVs were summarized into 5 groups “CF pathogens”, “strict anaerobes”, “facultative anaerobes”, “strict aerobes” and “unknown” according to the oxygen requirements of the respective taxon^19^. Next, the total data set was patient-stratified to avoid bias. 36 random data subsets with 650 samples each were generated by sampling with replacement from the total data collection. Subsequently, models with 1-25 DMMs were inferred stepwise for each subset using the R package *DirichletMultinomial.*^56^ Laplace approximation, BIC and AIC were queried independently to identify the optimal number of DMMs.

### Variance testing and ordination

To quantify the impact of covariates on the lung microbiome we applied PERMANOVA to rarified ASV data and the identical data remodeled by non-standard sample descriptors.^54^ Bray-Curtis distance was employed for ASVs and Euclidian distance for scaled and centered non-standard descriptors. The model was designed to test the marginal effects of the individual covariates (function *adonis2*, parameter setting by = margin). For comparison, we calculated effect sizes (ω^2^) of the covariates using the *adonis_omegaSq* function from the MicEco package.^57^ Principal component analysis was performed on scaled and centralized sample descriptors in R. Sample descriptors included Shannon diversity, Chao1 richness, relative abundance of the most abundant CF pathogen, the ratio of CF pathogen counts to counts from anaerobic taxa and the classification to a particular DMM community class.

### PEx type clustering and sample classification

Hierarchical k-mer clustering of samples was conducted on the first three principal components of the ordinated sample data using the *R package pheatmap.*^58^ Pearson correlation was employed as similarity measure. Next, χ^2^ contingency test was used to identify the best k cluster number for the classified sputum samples. Subsequently entire time series and networks were assigned to a k-mer cluster by majority vote of the included samples. Detailed inclusion criteria are explained in tables S1, S2.

### Statistical modeling of time behavior

Linear mixed effect models to determine time dependency of Shannon diversity and Chao1 richness were built using time groups (< 24;24-60 days before treatment; PEx clusters *PAT, AN1*) or 5 days intervals (PEx cluster 3, *AN2*) as fixed effects and subject ID, age group as well as PEx cycle id as random effects (package *lmerTest).*^59^ Standardized effect sizes (ω^2^) of predictors were calculated using the package *effectsize.*^60^

ANCOVA models were implemented using time groups (< 24, 24-60 days) as categorical predictors, sampling distance as numerical covariate and Aitchison distance as dependent variable. Aitchison distances were calculated using R package *robCompositions.*^61^

Graphical representations of boxplots and regression models were generated using *ggplot2*^62^ and *ggpubr*^63^.

### Co-occurrence network inference and network statistics

Co-occurrence networks were inferred from 20 samples collected at subsequent days with *SparCC* in a sliding window along each time series.^64^ Missing samples were imputed using R package *seqtim*e^32^ and jump size for the sample window was set to 1.

For downstream analysis on giant network components, only strong (ρ > |0.2|) and statistically significant (p < 0.01 after FDR correction) co-occurrence edges were included, as well as networks with < 10 imputed samples. Topological properties were assessed using R package *igraph.*^65^

Node degree and node clustering distributions were calculated across all networks classified in the same PEx type. To identify power laws and their respective slopes, linear regressions were performed on log-transformed data using R package *ggplot2.*^62^

For comparing the effect size of PEx clusters with covariates Shannon and Chao1 on graph topology, we first calculated clustering coefficient, betweenness centrality, node counts and edge counts for each giant component and averaged Shannon and Chao1 of all samples included for inference of the respective network. Next, we implemented independent LMMs, corrected for subject id and assessed the effects sizes (partial η^2^) as described previously.

To quantify the presence of certain ASVs in top hierarchical network positions, we first selected positions with relative highest degree (> 90% of all degree values) and relative lowest clustering (< 10% of all clustering coefficients). Subsequently, the frequency of ASVs on these positions was counted, normalized and ranked. All calculations were performed in R.

### Noise analysis of ASV time behavior

For each time series, noise colors of participating ASVs were inferred using package *seqtime* as previously described.^32^ In short, missing samples were interpolated, and rounded to counts, negative interpolation values were set to 0 counts. Subsequently, the wrapper function *identifyNoisetypes()* performed a spectral density estimate and calculated a linear fit to the resulting periodogram (log frequency vs log spectral density) of the time series. According to the slope of the fit, time series were classified into categorical noise color groups. Noise colors were plotted as ratios of pathogens vs anaerobic ASVs with similar color (relative abundances) in the same sample.

### Pathogen removal

To identify the dominant pathogen by network, the subset of samples used to infer the individual co-occurrence network was queried and the pathogen with highest cumulative abundance was selected. Only networks with dominant pathogen locating to the major component were used for further analysis. We calculated modularity, the number of unconnected components and node size of the giant component independently for each co-occurrence network before and after pathogen removal. All parameters were normalized to the corresponding value before node removal. Density distributions of the normalized parameters were scaled and plotted with *ggplot2* function *geom_density().*^62^ To test for difference of the cumulative parameter distributions, two-sided Kolmogorov-Smirnov tests were performed (*ks.test()*, *stats* package).^66^

## Author contributions

SW designed the overall study and analyses. JJL and SW engaged in acquisition of the financial support for the study leading to this publication. SW and KO performed the data analyses. JJL, LAC, LJC, LMK planned the daily sputum study, coordinated with participants, and collected samples, meta- and 16S sequencing data. GB provided important input on graph theory applied in this study. SW, JJL, KO, LAC, LJC participated in discussions related to this work. SW and JJL wrote the manuscript. All authors read and approved the final manuscript.

## Acknowledgements

We gratefully acknowledge the individuals who provided the sputum samples used in this study. We thank LiPuma lab staff for processing samples. This work was supported by National Institutes of Health grants R01HL136647 and R56HL126756 and Cystic Fibrosis Foundation grants LIPUMA13I0 and LIPUMA15P0 (JJL). SW was supported by the Austrian Science Fund (FWF) Elise Richter project V585-B31. We thank Ginestra Bianco for critical discussion of our network models and Philipp Starkl for valuable feedback on figure design.

## Data and code availability

The generated sequencing data (FASTq files) and associated meta-information are available as NCBI BioProject PRJNA987026, NCBI BioProject PRJNA520924.

