## Supplemental figures and tables for "Microbial community organization designates distinct pulmonary exacerbation types and predicts treatment outcome in cystic fibrosis"

### Supplement

#### Figure S1

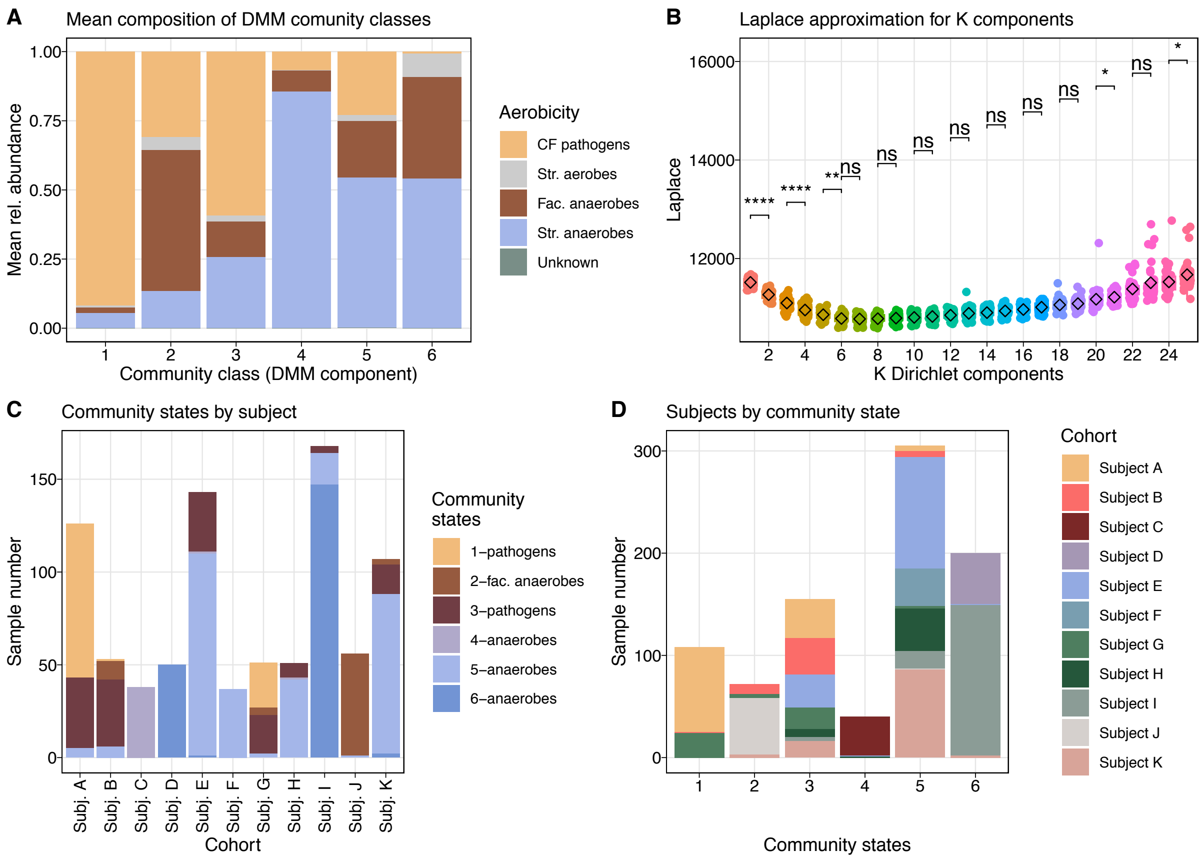


**Figure S1. Identification of microbial community classes.** ***(A)*** *Composition of 6 DMM components. DMM was inferred at the level of aerobicity class ("conventional CF pathogens", "strict aerobes", "facultative anaerobes", "strict anaerobes", "unknown").* ***(B)*** *Identification of best DMM model. 36 models, each with 1 to 25 components, were inferred and their Laplace approximations are presented together with stepwise comparison of means among models with k components (Wilcoxon). The first statistically non-significant change of Laplace was used as a cutoff criterion (k=6).* ***(C)*** *Distribution of DMM community states across cohort. Sample number by subject is depicted.* ***(D)*** *Distribution of subjects across DMM components. Sample number by DMM is depicted.*

#### Figure S2

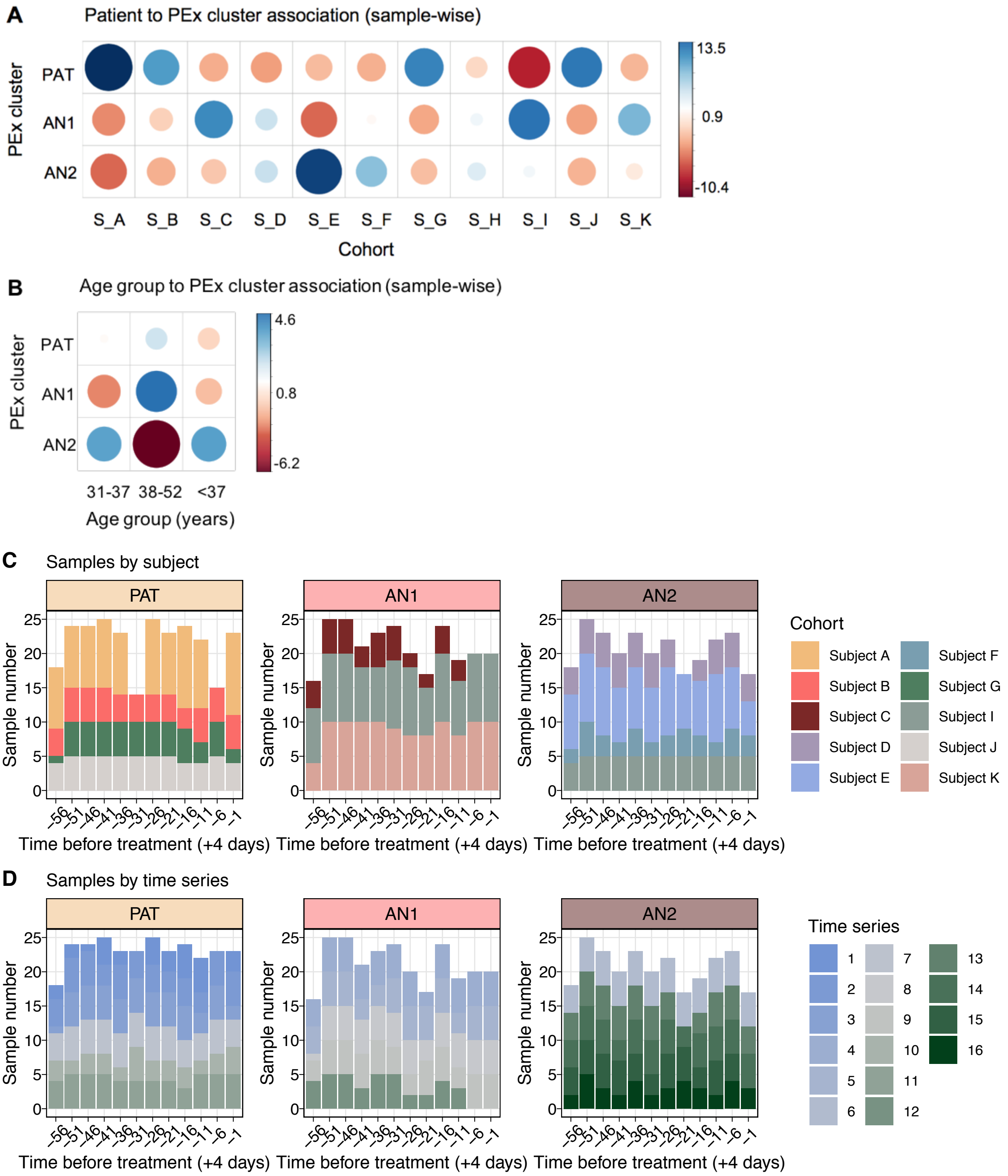


**Figure S2. Sample distributions by PEx clusters, subjects, age groups and time series.** ***(A)*** *Spearman rank association of PEx clusters and subjects.* $\chi^2$ *testing performed on samples in respective categories, standardized residuals shown* $(\chi^2 = 700,\ p < 2.2e-16)$. ***(B)*** *Spearman rank association of PEx clusters and age groups.* $\chi^2$ *testing performed on samples in respective categories, standardized residuals shown* $(\chi^2 = 42.5, p < 1.3e-8)$. ***(C)*** *Sample distribution by subjects and*

*time to acute treatment as used in downstream analyses.* ***(D)*** *Sample distribution by time series and time to acute treatment as used in downstream analyses.*

#### Figure S3

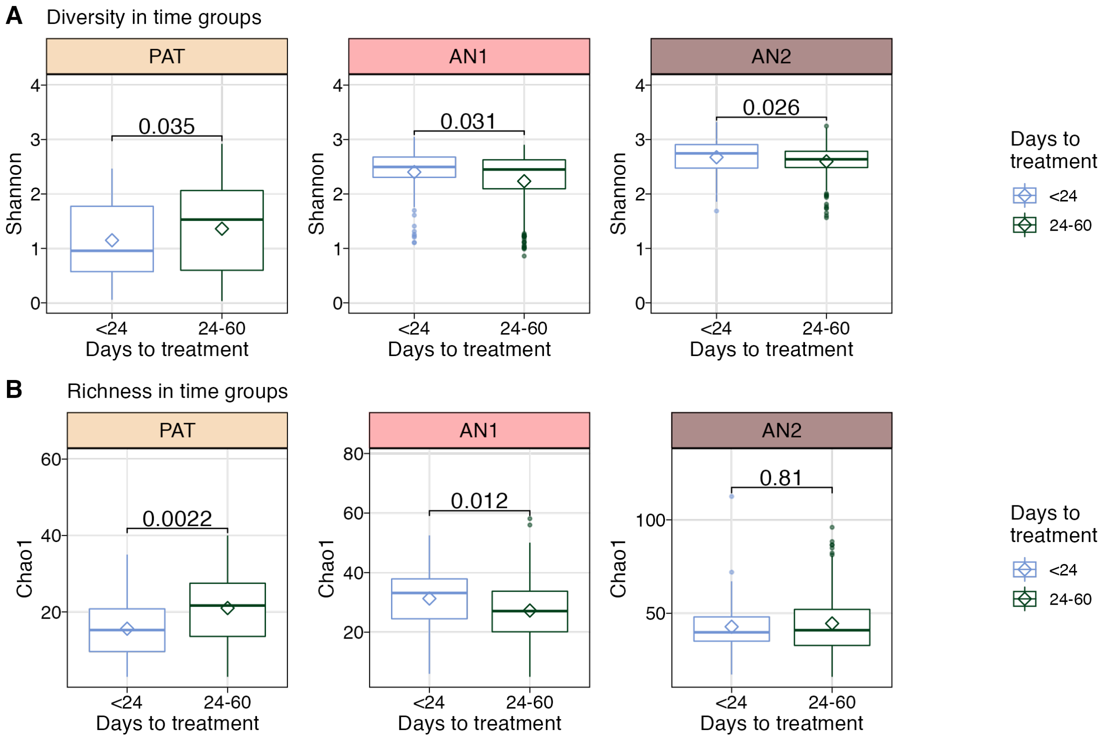


**Figure S3. Time evolution of microbiome diversity and richness by time group and PEx cluster.** ***(A)*** *Shannon diversity in microbiomes collected either <24 days or 24-60 days before acute treatment. Significance (p value) of group-wise Wilcoxon tests depicted.* ***(B)*** *Chao1 richness in microbiomes collected either <24 days or 24-60 days before acute treatment. Significance (p value) of group-wise Wilcoxon tests depicted.*

#### Figure S4

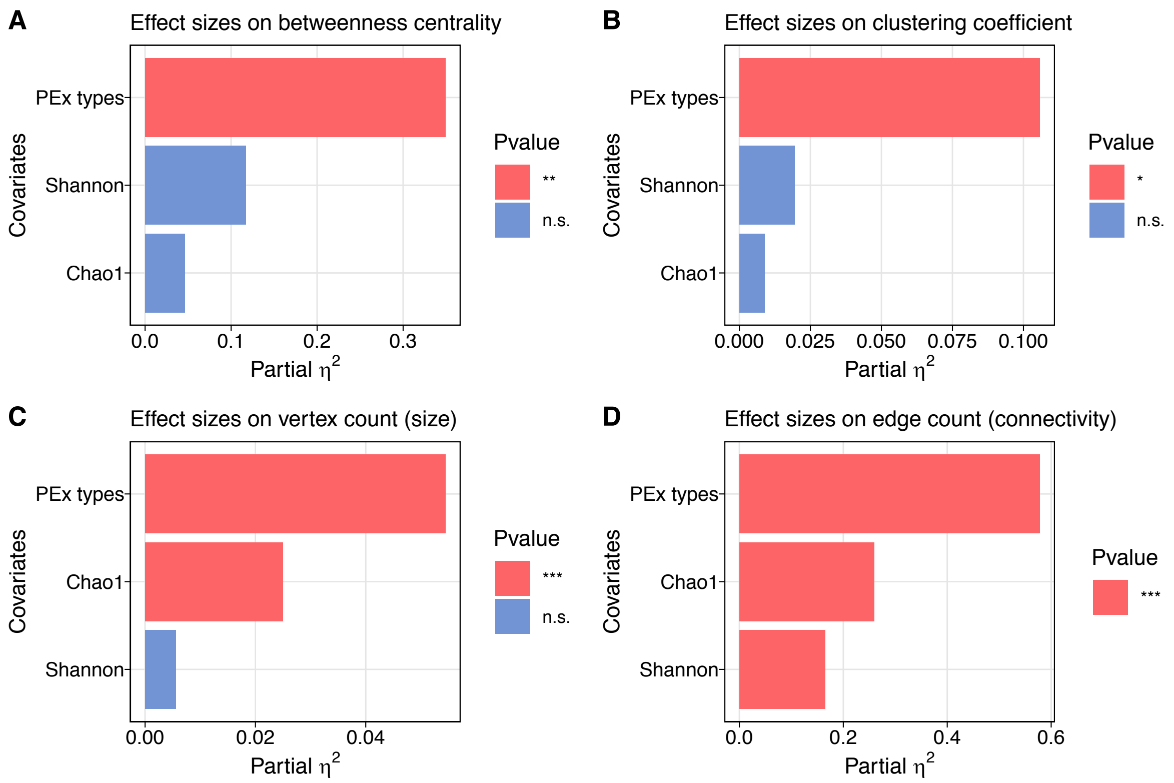


**Figure S4. Effect sizes of PEx clusters, and covariates Shannon diversity and Chao1 richness on co-occurrence topology.** ***(A-D)*** *Independent mixed effect models were implemented and the partial effect sizes, as well as p values, of covariates on betweenness centralities* ***(A)****, clustering coefficients* ***(B)****, vertex counts* ***(C)*** *and edge counts* ***(D)****, were assessed and visualized. Solely giant components of co-occurrence networks were included. Levels of statistical significance displayed by color code in legend.*

**Table S1 Inclusion criteria for time series & samples together with performed analyses.**

| Inclusion criteria | Samples | Subjects | Time series | Samples by subject mean (sd) | Samples by time series mean (sd) | Time series by subject mean (sd) | Analyses |
|---|---|---|---|---|---|---|---|
| >=60 days w/o acute treatment | 880 | 11 | 18 | 80.0 (46.9) | 48.9 (7.3) | 1.6 (0.9) | DMM<br>PCA<br>clustering<br>NW inference<br>noise colors |
| >=60 days w/o acute treatment<br>>=40% time series coverage by PEx cluster | 789 | 10 | 16 | 78.9 (44.4) | 49.3 (7.4) | 1.6 (0.8) | time series classification<br>temporal evolution<br>community turnover |

**Table S2 Inclusion criteria networks together with performed analyses.**

| Inclusion criteria | NWs | Subjects | NWs by PEx cluster PAT; AN1; AN2 | NWs by subject mean (sd) | Analyses |
|---|---|---|---|---|---|
| Samples from classified time series<br>NW classification by majority vote of sample cluster association | 589 | 10 | 222; 192; 175 | 58.9 (31.3) | NW classification<br>topological analyses<br>taxonomic hierarchy<br>treatment simulation |

**Table S3 Kolmogorov Smirnov test to contrast cumulative distributions of network properties before and after simulated pathogen removal.**

| Kolmogorov Smirnov distribution test | | | |
|---|---|---|---|
| Network property | Test cases | D statistic | p-value |
| No. nodes | PAT vs AN1 | 0.33 | 2.7e-04 |
| | PAT vs AN2 | 0.38 | 3.8e-08 |
| Modularity | PAT vs AN1 | 0.30 | 2.7e-03 |
| | PAT vs AN2 | 0.45 | 5.4e-10 |
| Strong components | PAT vs AN1 | 0.56 | 1.5e-11 |
| | PAT vs AN2 | 0.49 | 1.1e-13 |
